# Surface interaction patches link non-specific binding and phase separation of antibodies

**DOI:** 10.1101/2022.03.07.483238

**Authors:** Hannes Ausserwöger, Georg Krainer, Timothy J. Welsh, Tomas Sneideris, Matthias M. Schneider, Gaetano Invernizzi, Therese W. Herling, Nikolai Lorenzen, Tuomas P. J. Knowles

## Abstract

Non-specificity is a key challenge in the successful development of therapeutic antibodies. The tendency for non-specific binding in antibodies is often difficult to reduce via judicious design and, instead, it is necessary to rely on comprehensive screening campaigns. A better understanding of the molecular origins that drive antibody non-specificity is therefore highly desirable in order to prevent non-specific off-target binding. Here, we perform a systematic analysis of the impact of surface patch properties on antibody non-specificity using a designer antibody library as a model system and DNA as a non-specificity ligand. Using an in solution microfluidics approach, we discover patches of surface hydrogen bonding to be causative of the observed non-specificity under physiological salt conditions and suggest them to be a vital addition to the molecular origins of non-specificity. Moreover, we find that a change in formulation conditions leads to DNA-induced antibody liquid–liquid phase separation as a manifestation of antibody non-specificity. We show that this behaviour is driven by a cooperative electrostatic network assembly mechanism enabled by mutations that yield a positively charged surface patch. Together, our study provides a direct link between molecular binding events and macroscopic liquid–liquid phase separation. These findings highlight a delicate balance between surface interaction patches and their crucial role in conferring antibody non-specificity.

## Introduction

Antibodies have emerged as highly potent drug molecules, particularly due to their high specificity, long *in-vivo* half–life, and ability to induce immunogenic effector functions^1,2^. As of 2021, more than hundred therapeutic antibodies for the treatment of various human diseases, including cancer^3–5^, Alzheimer’s disease^6^, osteoporosis^7,8^, and HIV^9^, have been approved by the United States Food and Drug Administration^10^. Despite the enormous success of antibody-based pharmaceuticals, only 21 % of candidates admitted to clinical trials have been approved^11^. This adds to significant developmental costs of 1.4 billion Dollars per approved compound^12^. The primary reasons for these failures are likely due to unsatisfactory clinical readouts and commercial considerations. Interestingly, however, it has been reported that antibodies on the market posses lower tendency for non-specificity compared to candidates that fail in clinical trial Phase 2 or 3,^13,14^ suggesting that unsuccessful clinical trials are associated with undesired non-specific binding.

Indeed, target specificity is often hard to achieve in the development of therapeutic antibodies, and as a result, antibodies that exhibit non-specific off-target binding are commonly generated (Figure 1). This is because target specificity is not an inherent property of antibodies^14–20^, as replication of the meticulous *in vivo* maturation process is very difficult to obtain with similar precision *in vitro* due to a trade-off between affinity and specificity^17,21^. Nonetheless, obtaining highly specific antibodies is crucial as several recent reports have demonstrated how insufficient specificity often manifests in critical development issues (Figure 1b), such as decreased therapeutic effects due to sub-optimal pharmacokinetics or fast *in vivo* clearance rates^22–28^ or even *in vivo* toxicity.^29^ Moreover, non-specificity has also commonly been linked with physical stability issues related to bioprocessing and formulation such as high solution viscosity, low solubility and phase separation^26,28,30–32^. Hence, specificity is not only the strength of antibodies but in many ways also their weakness and a key hindrance to their developability (i.e., the overall suitability for successful development into a drug).

**Figure 1:**
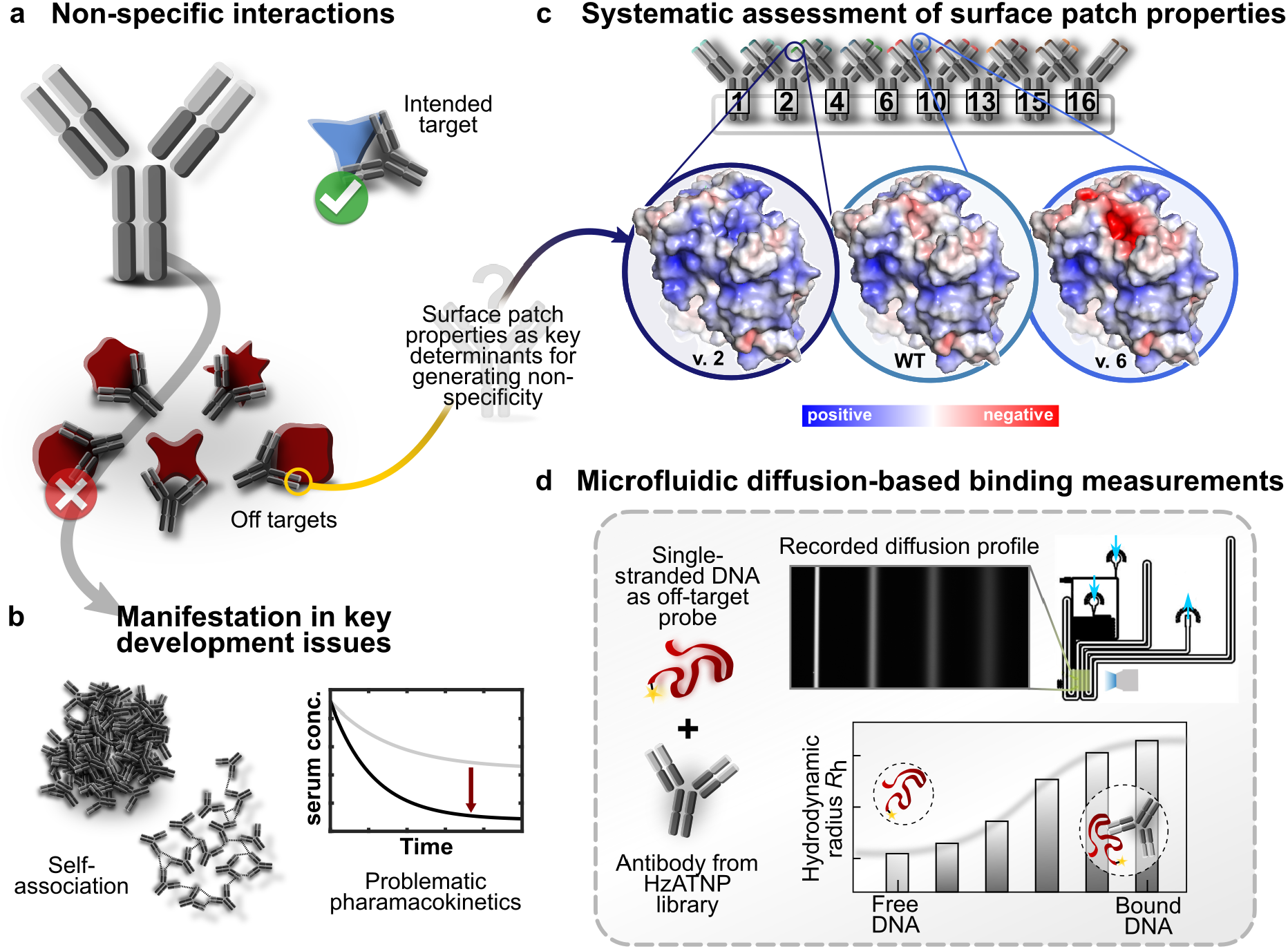
Probing antibody non-specificity in a rationally designed antibody library using an in solution microfluidics approach. **(a)** Non-specificity is defined as the binding of antibodies to unwanted target binding partners. **(b)** This commonly leads to problematic manifestations in important development issues such as self–association^26,28,30,31^ and undesired *in vivo* pharmacokinetics^22–29^. Individual case studies have highlighted surface patches properties, i.e. the extent of groupings of amino acids with similar physico-chemical properties on the protein surface, as key contributors to non-specificity^28^. **(c)** In our study, we perform a systematic assessment of the impact of surface patch properties on non-specific binding using the HzATNP library, which was generated to represent differences between surface property distributions such as the surface charge potential. **(d)** Non-specific binding is probed against single-stranded DNA oligos and monitored using microfluidic diffusional sizing (MDS). MDS measures the diffusion profiles of labelled molecular species in a microfluidic chip, to determine their hydrodynamic radii. Complex formation, entailed by non-specific binding events, yields an increase in the observed hydrodynamic radius allowing for quantitative solution phase study of non-specific antibody–DNA binding.

Despite being recognised as a vital developmental issue for the generation of therapeutic antibodies^13,33^, the molecular causes of non-specificity on the protein level remain largely unaddressed. As such, it is unclear which molecular features drive non-specificity and how they mechanistically lead to the aforementioned problematic macroscopic manifestations. Furthermore, it remains to be discovered if and how formulation and environmental conditions impact non-specificity behaviour. Individual case studies investigating the importance of surface patches^25,26,28,31,34^ (i.e., groupings of amino acids with similar physico-chemical properties on the protein surface) have shown high potential in addressing non-specificity behaviour. One example is the work of Dobson *et al*.^28^ who studied the role of a hydrophobic amino acid patch on the protein surface of the MEDI1912 antibody. The surface patch leads to extensive non-specific binding which translates into critical development issues like high *in vivo* clearance rates and high solution viscosity, all of which could be removed by mutational disruption. Such considerations of surface patches remain a rarity and thus, surface patch properties are yet to be investigated more systematically using antibody libraries.

In addition, insights into the molecular origins of non-specificity have been hampered by methodological shortcomings. To date, the study of non-specific binding usually relies on determining interactions through surface immobilisation techniques or indirect measurements of physico-chemical properties. These approaches, however, fail to accurately represent binding events under native conditions (i.e., directly in solution without surface and avidity effects), which for weak, low affinity interactions characteristic for non-specificity can yield substantial changes in the perceived properties^35^. Approaches able to probe non-specificity effects in solution are therefore highly desirable.

Here we provide insights into the molecular origins of antibody non-specificity by examining non-specific binding in a rationally designed antibody library using an in solution microfluidics approaches. Specifically, we study the HzATNP antibody library, designed with CamSOL, to represent systematic variations of the physico-chemical surface patch properties^30,36^ (Figure 1c). This library represents an ideal model system for conceptual investigation as it has been characterized in detail to show distinct changes in the protein biophysical properties between the individual variants.^37^ We assess the impact of the changes between variants on the non-specific binding to single-stranded DNA, a common non-specificity ligand which is a valuable indicator for the overall propensity of non-specific interactions also *in vivo*^27,33,38^. By establishing microfluidic diffusional sizing (MDS)^39,40^ (Figure 1d) as a tool for probing antibody non-specificity, we identify that non-specific binding of HzATNP antibodies to DNA at physiological salt conditions is driven by a patch of amino-acid residues that engage in hydrogen bonding. Indeed, only individual, rationally selected mutations modulating the surface hydrogen bonding properties can reduce the propensity for non-specific binding by decreasing DNA affinities by multiple orders of magnitude. Investigations under lowered salt conditions reveal that the formulation conditions have a crucial impact on the protein physico-chemical properties determining non-specificty behaviour. This leads to the observation of DNA-induced liquid–liquid phase separation (LLPS) of antibodies for a subset of the library as a macroscopic manifestation of non-specificity due to the culmination of many weak multivalent interactions.^41^ We further characterised the transition from individual molecule binding to the phase separated state via mapping the phase space using PhaseScan, a high-throughput droplet microfluidics based approach^42^ and combining it with detailed analysis of binding events. Together, our study provides first mechanistic links between microscopic causes and macroscopic manifestations of antibody non-specificity. In doing so our study highlights the delicacy of how individual mutations and formulation conditions can alter the balance between charged and hydrogen bonding patches in governing non-specificity binding and phase separation of antibodies.

## Results

### Probing the influence of surface patch properties on non-specific binding in the HzATNP antibody library by microfluidic diffusional sizing

The HzATNP library consists of a total of 17 antibodies designed around the humanized anti-trinitrophenyl wild type (WT) antibody, to represent a variety of protein solubilities. The library was generated by Wolf-Perez *et al*.^30,36^ using a computational design algorithm in CamSol, which predicts solubility based on the extent and the frequency of physico-chemical surface property hot spots. The library consists of variants with charged and/or hydrophobic mutations that modulate the protein surface patch properties by disrupting or enhancing aforementioned hot spots. As such, this antibody library represents rationally selected, systematic changes in the surface patch properties, making it an ideal model system to study non-specific binding. We investigated a subset of the library, consisting of the WT antibody as well as variants 1, 2, 4 and 6 (charged mutation variants) and variants 10, 13, 15 and 16 (hydrophobic mutation variants). The exact positions of the mutations in the HzATNP variants investigated here are summarised in Table 1. As a non-specificity ligand, we used single-stranded DNA, which is commonly used as a proxy to probe the overall propensity of antibody non-specificity,^14^ likely by probing potential charge and nucleotide hydrogen bonding interactions.

**Table 1:** Summary of the mutations in the Hz-ATNP antibody library. Mutations implemented in the HzATNP library around the wild type (WT) antibody in the heavy chain (HC) and light chain (LC) protein domains. Variants 1, 2, 4 and 6 are generated by introducing charged residues and variants 10, 13, 15 and 16 are obtained by mutagenesis to hydrophobic residues.

| Variant | HC Mutations | LC Mutations |
| --- | --- | --- |
| 1 | T68G, Q108E | T5D, V99R |
| 2 | T68G, Q108E | V99R |
| 4 | T68A | V99K |
| 6 | S70E | V99D |
| WT | - | - |
| 10 | - | K193V |
| 13 | E16V, K120V | - |
| 15 | E16V, K120F | K193Y |
| 16 | E16F, D72F, K120W | K193F |

To measure the interactions between antibodies and DNA, we adopted the in solution microfluidic-based approach MDS^39^ to acquire diffusion profiles of molecular species within a microfluidic channel. From broadening of the diffused species in space and time the diffusion coefficient *D* can be extracted and the corresponding hydrodynamic radii *R*_h_ can be calculated using considerations of the advection-diffusion process and the Stokes–Einstein relation^39,40^. Binding events can be observed upon complexation of binding partner to the labelled species as it leads to an apparent increase in the observed hydrodynamic radius. Using MDS, we measured the hydrodynamic radii of Cy3-labelled single-stranded DNA (50 mer of random sequence, see methods section) at a constant concentration of 1 µM in the presence of an excess of the individual antibody variant (6.7 µM) (Figure 2). Interactions were probed under physiological salt conditions (20 mM HEPES buffer pH = 7.4, 150 mM NaCl, supplemented with 0.01% (*v/v*) Tween). The excess in antibody concentration was chosen to ensure binding saturation or, at least, observe a significant size increase for the DNA oligomer. We also performed experiments, in which we varied the antibody concentration for selected variants over several orders of magnitude to obtain binding curves and determine the apparent affinity of the antibody–DNA interaction (see Figure 3).

**Figure 2:**
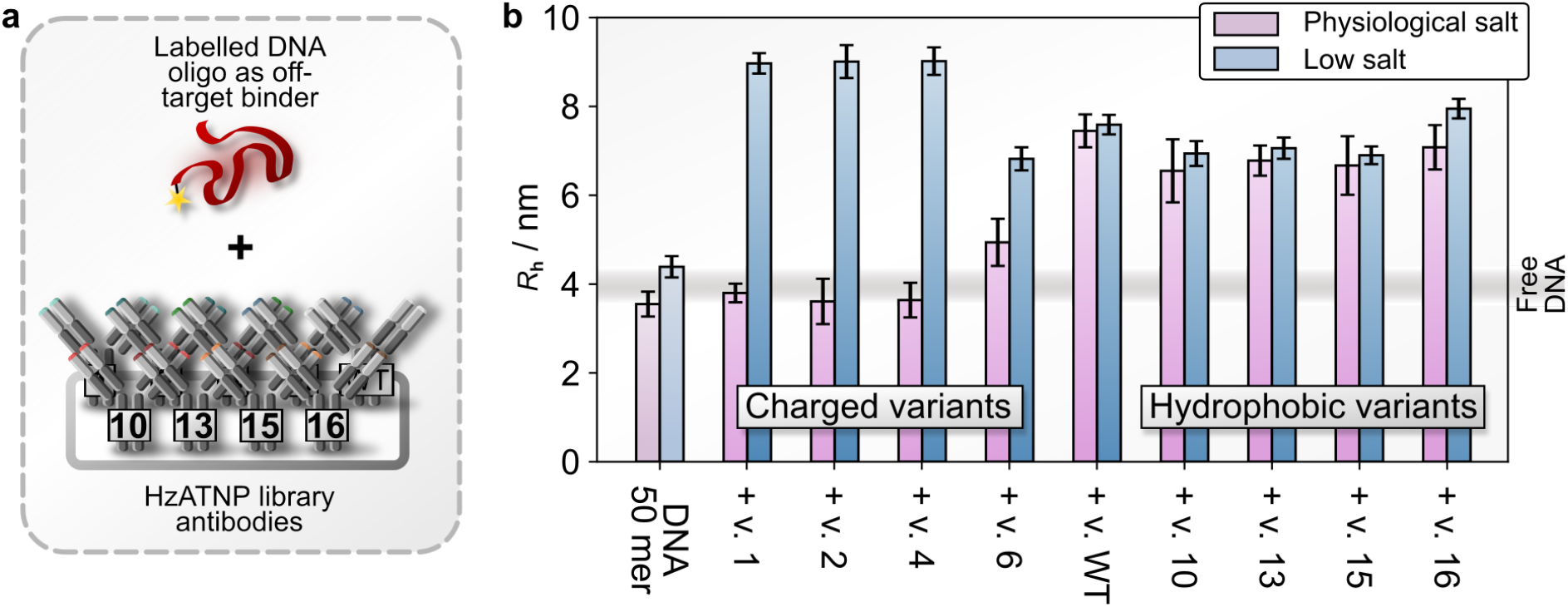
Hydrodynamic radius changes of labelled DNA oligos in complexation with HzATNP library antibodies. **(a)** Single-stranded DNA 50 mer labelled with Cy3 is subjected to binding with the various HzATNP antibody library variants under physiological salt (20 mM HEPES buffer, pH = 7.4; 150 mM NaCl, with 0.01% (*v/v*) Tween) and low salt conditions (2 mM HEPES buffer, pH = 7.4; 15 mM NaCl, with 0.01% (*v/v*) Tween) at 1 µM DNA and 6.7 µM antibody concentration, after which the hydrodynamic radii (*R*_h_) were recorded via microfluidic diffusional sizing (MDS). **(b)** At physiological salt conditions the WT as well as variants with hydrophobic mutations (variants 10, 13, 15 and 16) display binding which is largely unchanged by decrease in ionic strength. For variant 6 at physiological salt conditions, a decrease in the hydrodynamic radius, indicative of a decrease in affinity, is found. Variants 1, 2 and 4 display strongly decreased tendency to bind non-specifically at physiological salt conditions whereas large complexes were formed with DNA at lowered ionic strengths. Errors represent standard deviations from triplicate measurements.

**Figure 3:**
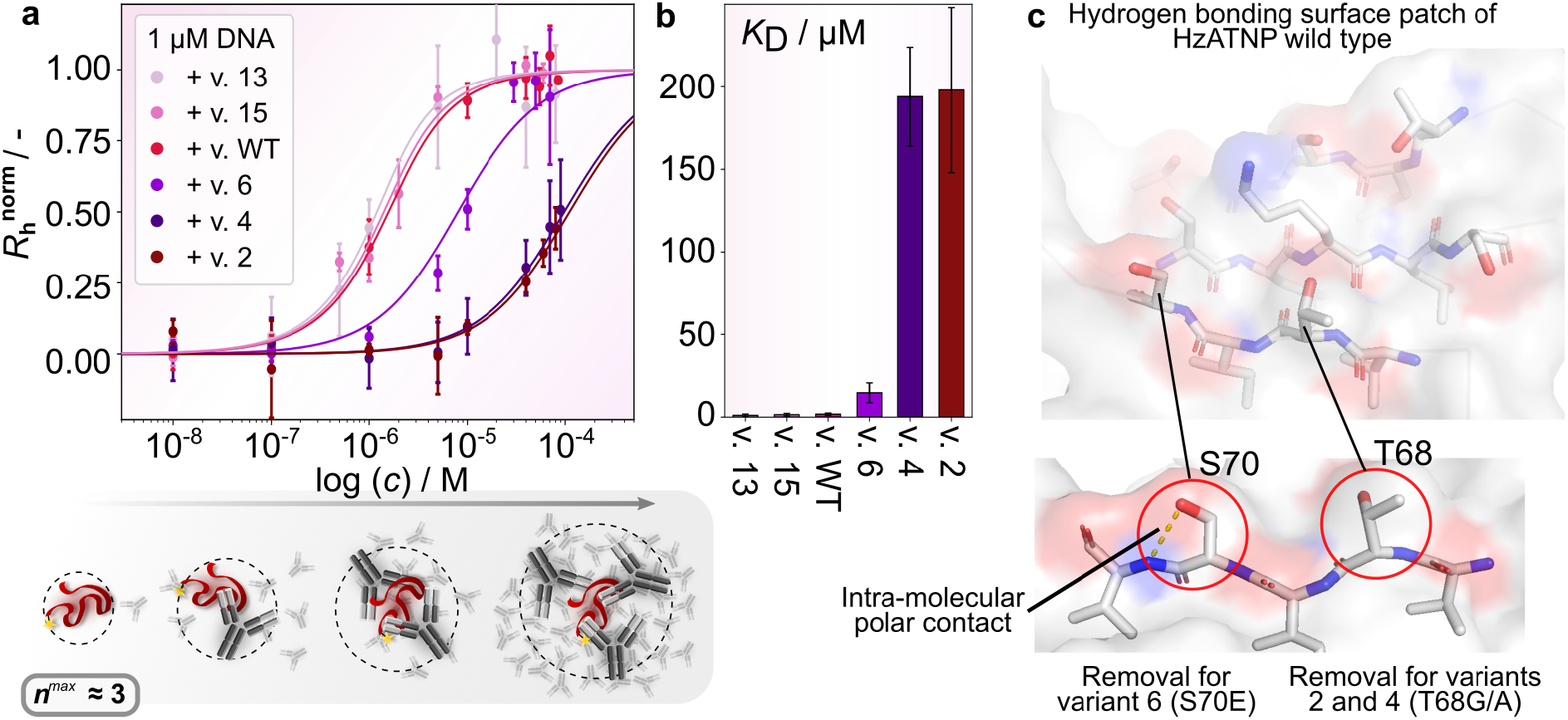
Non-specificity binding curves of selected HzATNP antibody variants against DNA and investigation of the hydrogen bonding surface patch. **(a)** Changes in the normalized hydrodynamic radius of DNA at 1 µM with varying antibody concentrations of selected variants at physiological salt conditions (20 mM HEPES buffer, pH = 7.4; 150 mM NaCl; supplemented with 0.01% (*v/v*) Tween). Non-normalized *R*_h_ values range from ≈3.5-7 nm. Errors represent standard deviations from triplicate measurements. At high excesses of antibody concentration, binding site saturation on the DNA is observed, hence, plateauing of the hydrodynamic radii. Between variants, a gradual decrease in binding affinity from the hydrophobic variants (represented by variants 13, 15 and WT) to variant 6 and further the charged variants (represented by variants 2 and 4) becomes apparent. Stochiometric analysis of the antibody–DNA complexes on the WT reveals that ≈ 3 antibody molecules bind to individual DNA strands until DNA binding site saturation is reached. **(b)** Comparison of the dissociation constant *K*_D_ of the interaction of different variants to DNA, highlighting drastic changes in affinity with only individual mutations. **(c)** Molecular model of the protein surface in the vicinity of residues crucial to non-specific binding (T68, S70). Oxygen and Nitrogen atoms are coloured red and blue respectively, to indicate points with a tendency to engage in polar contacts. All other atoms are shaded in grey. This shows multiple residues able to engage in hydrogen bonds (T17, S19, T21, T68, S70, S79 and K81) clustered on the protein surface, hence, likely forming a hydrogen bonding surface patch responsible for the non-specific binding to DNA of HzATNP WT antibody. Additionally, a close-up view of both relevant residues for the HzATNP antibody library (S70 and T68) is provided showing an intra-molecular polar contact of S70 to the amide backbone and the free side chain of T68.

### Hydrogen bonding surface patches as potential key determinants driving non-specificity

First, we probed potential complexation of the HzATNP antibody variants with DNA based on non-specific interactions. Indeed, upon mixing of the WT antibody with the DNA an increase of the DNA hydrodynamic radius from 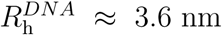 for free DNA 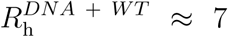 was observed. We further examined the binding affinity of the non-specific interactions between the WT antibody and DNA (Figure 3a) by recording a binding curve and obtained a *K*_D_ value in the micromolar regime 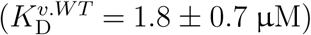 (Figure 3b). From a mechanistic perspective, the observed non-specific binding is either driven by electrostatic interactions to the highly negatively charged nucleic acid backbone or coordinated hydrogen bonding interactions with the available nucleobases/phosphates as well as potential *π*-stacking and hydrophobic interactions between nucleobases and compatible amino-acid residues.

Next, we probed the interaction of DNA with antibody variants of the hydrophobic series (variants 10, 13, 15 and 16). These variants (Figure 2b) did not exhibit a significant change in behaviour compared to the WT antibody, as the size of the detected complexes remained constant, ≈ 7 nm. This suggests that additional hydrophobicity does not increase the propensity of the antibodies to bind DNA non-specifically. We further measured the binding affinity of selected hydrophobic variants (Figure 3a) and found that there is no apparent difference in DNA non-specific binding between the WT antibody and the hydrophobic variants (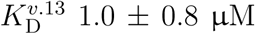 and 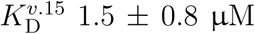 vs. 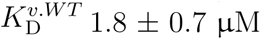) (Figure 3b). This corroborates that hydrophobic interactions are unlikely causative for the observed non-specificity, as the changes in hydrophobicity implemented in the hydrophobic variants have no noteworthy impact on it.

Finally, we probed the charged variant series (variants 1, 2, 4 and 6). These exhibited a reduced propensity to form complexes with DNA compared to the WT antibody (Figure 2b). At 1 µM DNA and 6.7 µM antibody concentration, only variant 6 exhibited binding to DNA 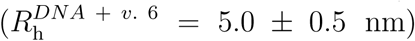, however, at a reduced complex size compared to the WT antibody. More specifically, this corresponds to about ∼60% binding DNA binding site saturation. Variant 2 and 4 displayed no binding. The decrease in binding appears to be independent of the overall net charge change introduced by the mutations 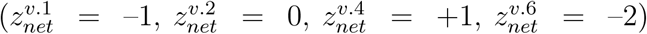. Hence, the observed non-specificity is very unlikely to be electrostatically driven. This suggests that hydrogen bonding is likely the dominant contributor to the observed non-specificity, as neither hydrophobic nor electrostatic variations appear to have a pronounced impact. Indeed, the binding to DNA appears to correlate with the presence (or absence of) amino acids that can engage in hydrogen bonding. Analysis of binding affinities confirms that variants, in which amino-acid residues that can engage in hydrogen bonding were removed (variant 1, 2, 4 and 6), display a decreased binding affinity to DNA (Figure 3a, b). Compared to the WT antibody 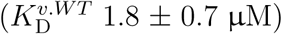, we find an affinity of 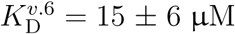 for variant 6 which carries the S70E and V99D mutations. A further reduction to 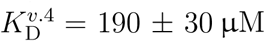 and 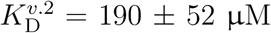 is observed for variants 4 and 2, respectively, which have the T68G/A and V99K/R mutations common also with variant 1. As a result, the reduction in affinity from the WT antibody to variant 6 and variants 2 and 4 is likely a result of decreased hydrogen bonding engagement. This is also represented in the reduction of affinity from variant 6 to variants, 2 and 4 based on the nature of the mutations (S70E on variant 6 vs. T68G/A on variants, 2 and 4). Glutamic acid as a charged amino-acid residue is likely to replicate some polar contacts engaged by the serine hydroxy functional group, whereas Glycine, as a small, non-polar side chain, is not, explaining the tighter binding of variant 6 over variant 2 and 4.

To further investigate the role of hydrogen bonding in the non-specific binding of HzATNP antibody variants, we took advantage for the homology model structure generated of the WT antibody.^30^ As shown in Figure 3c, the crucial residues T68 and S70 are spatially surrounded on the protein surface by multiple other residues that are able to engage in hydrogen bonding (T17, S19, T21, S79 and K81), likely allowing for concerted hydrogen bond complexation. Additionally, S70 is engaged in an intra-molecular polar contact with the amide backbone, whereas T68 is not, which can further explain the drop in DNA affinity from variant 6 (S70E) to variants 1, 2 and 4 (T68G/A). Crucially, this suggests that non-specific binding can arise by clusterings of residues able to engage in hydrogen bonding on the protein surface. This, to the best of our knowledge, is the first description of hydrogen bonding surface patches in non-specific binding to DNA, and is therefore a key addition to the discussion around the molecular origins of non-specificity.

### Reduction of ionic strength reveals DNA-induced phase separation of antibodies

In a next set of experiments, we performed MDS measurements at lowered ionic strengths to explore the role of electrostatic interactions in the observed non-specificity. Electrostatic contributions are expected to become increasingly dominant at lower ionic strengths, given the accompanying decrease in Debye screening. Moreover, varying ionic strength is a common strategy in protein and antibody development to investigate the impact of fundamental physico-chemical forces underlying association processes and to optimize formulation strategies^43–45^. To this end, we studied the interactions between Hz-ATNP library antibodies with DNA in buffer with reduced salt content (2 mM HEPES buffer pH = 7.4, 15 mM NaCl, with 0.01% (*v/v*) Tween).

We first probed interactions at 1 µM DNA and 6.7 µM antibody for all variants. As shown in Figure 2b, we find a size increase for pure DNA in low salt conditions as compared to physiological salt conditions (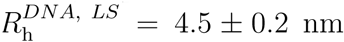 vs. 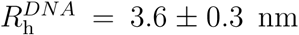). The size increase at low salt can be rationalized by an increase in monomer repulsion within the polymer chain^46^. For the interaction between the antibodies and the DNA, we observed limited changes to 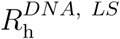 at lowered salt conditions for the WT antibody, the hydrophobic variants (variants 10, 13, 15 and 16) and variant 6 (Figure 2b). Drastic hydrodynamic radius increases, however, were obtained for the other variants of the charged mutation series (variants 1, 2 and 4; see Figure 2b). Variants 1, 2 and 4, all of which have increased positive charge and a reduced hydrogen-bonding surface patch, form large complexes up to ∼ 9–10 nm in hydrodynamic radius at low salt concentrations. We further performed control experiments using diffusional sizing in which one of the two components was left out of the solution mixture and found that neither the DNA nor the antibody had aggregated on their own due to low salt to drive formation of these large complexes (see Supplementary Figure S1). This indicates a change in the binding mode towards a non-stoichiometrically constrained electrostatic network assembly process. From stoichiometric analysis, we infer that complexes of 10 nm in size would require ∼10 antibody and ∼3 DNA molecules to come together (see Supplementary Information, ‘Affinity and stoichiometry analysis’), assuming a globular regime. Hence, we further refer to these complexes as high molecular weight clusters. By comparison, the stoichiometry of WT antibody–DNA complexes at physiological salt conditions was found to be ≈ 3 antibodies per an individual DNA strand for the binding site saturated state, as schematically represented in Figure 3a (detailed analysis in the Supplementary Information, ‘Affinity and stoichiometry analysis’ and Figure S2).

To explore further the formation of the high molecular weight antibody–DNA clusters, we performed binding titration experiments with a subset of antibodies (variants WT, 2 and 6) at lowered ionic strengths (Figure 4a). While variant 6 showed a typical binding curve with saturation, the WT antibody and variant 2 did not show binding saturation. A similar behaviour was determined for variant 4 (see Supplementary Figure S3). Although the sizes determined for the WT antibody are smaller compared to variant 2 at similar conditions (e.g. at 1 µM DNA and 6.7 µM antibody: 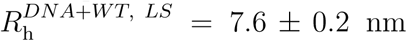 vs. 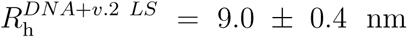, the hydrodynamic radius displayed continuous growth with increasing antibody concentration for both systems (Figure 4a). This suggests that binding of additional antibodies, in the case for the WT and variant 2 antibodies, can allow for recruitment of new DNA strands based on additive electrostatic interactions, as opposed to saturation of defined binding sites. Despite these variants being weaker binders at physiological salt conditions, they seem to have an increased propensity to form many weak interactions resulting in network assembly. The discriminator between these behaviours is likely the electrostatic repulsion between antibody and DNA within complexes. The behaviour of variant 6 particularly supports this as it carries multiple additional negative charges (i.e., S70E, V99D) and displays disruption of the electrostatic network assembly process given that no high molecular weight cluster formation was found. The binding observed, which follows a binding site saturation regime, is likely from residual hydrogen bonding observed under physiological salt conditions. Taken together, this indicates that mutations modulating the surface patch properties do not only impact molecular binding events but can even translate to macroscopic behaviours.

**Figure 4:**
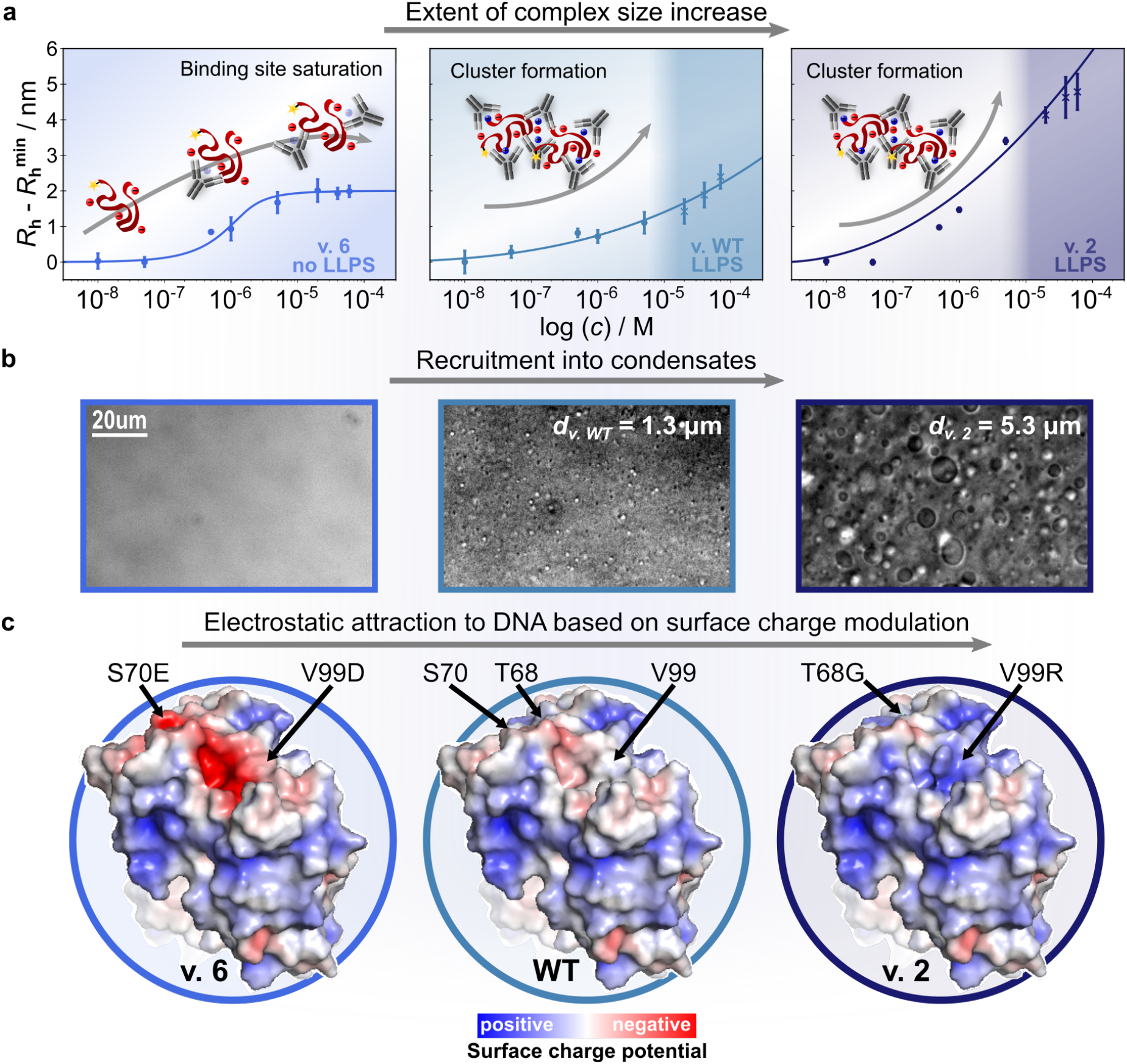
Changes in binding regime at lower ionic strengths reveal DNA-induced phase separation of antibodies. **(a)** Binding curve titration of selected HzATNP antibody variants against DNA at 1 µM and lowered ionic strengths (2 mM HEPES buffer, pH = 7.4; 15 mM NaCl; supplemented with 0.01% (*v/v*) Tween; 4°C). Variant 6 displays plateauing of the hydrodynamic radius due to bindingsite saturation as observed at physiological salt conditions. For the WT and variant 4 antibody, plateauing is not observed as complex sizes continuously increase with increasing antibody concentration, hence, indicating a change in binding regime towards electrostatic network assembly into clusters. This further leads to nucleation of condensates into a phase separated state. Points beyond the phase boundary were recorded in the supernatant post centrifugation. Errors represent standard deviations from triplicate measurements. **(b)** Brightfiled microscopy images of the observed macroscopic phase behaviour, recorded at at 40 µM antibody and 5 µM DNA. Variant 2 compared to the WT antibody displayed a more pronounced increase in complex size with higher antibody contents well as a significantly larger mean condensate size *d*. **(c)** Molecular models of the surface charge potential (PyMol, ABPS) highlighting the locations of residues subjected to mutation. This shows a large positively charged region in variant 2 likely allowing for additive electrostatic interactions to DNA. Disruption of this patch via removal of positive charges (WT) and introduction of negative charges (variant 6) creates additional electrostatic repulsion in DNA–antibody complexes, hence observed changes in binding behaviour.

The indication that some HzATNP antibody variants in the presence of DNA can form high molecular weight clusters at low ionic strengths based on additive electrostatic interactions suggests that mesoscopic / macroscopic phenomena may be at play. To investigate this further, we imaged mixtures of the above tested subset of antibodies (variants WT, 2 and 6) with DNA using microscopy (5 µM DNA and 40 µM antibody) (Figure 4b). Strikingly, for the WT antibody and variant 2, we observed spherically shaped assemblies that readily fused and merged. This indicates that these assemblies are phase separated condensates resulting from a demixing of the antibody and DNA through liquid-liquid phase separation. Control experiments with both DNA and antibody at the same conditions confirm that neither DNA nor antibody display this phase separation behaviour on their own (see Supplementary Figure S4). Further testing with 20 and 100 mer DNA as well as PolyA RNA (700–3500 kDa) also yielded phase separated condensates, suggesting that the behaviour is not based on an interaction specific to a particular secondary DNA structure or length (Supplementary Figure S5 and S6). Lastly, we identified that the demixed phase undergoes complete dissolution upon disruption of electrostatics via addition of high salt content but leads to aggregate formation upon addition of the hydrophobic disruptor in 1,6-hexanediol (see Supplementary Figure S7). This corroborates the initial suggestion that the observed behaviour represents phase separation of antibodies with DNA based on additive electrostatic interactions. Interestingly, when imaging directly after mixing, condensates formed by variant 2 tend to form larger condensates on average compared to WT under the same conditions (droplet diameter: *d*_*v*.2_ = 5.3 ±1.7 µm, *d*_*v*.*WT*_ = 1.3 ± 0.4 µm both at 40 µM antibody and 5 µM DNA) (Figure 4b). This more effective recruitment into condensates is an indication of less electrostatic repulsion within antibody–DNA complexes for variant 2.

To rationalize the impact of mutations on the electrostatically driven phase separation behaviour of antibody variants, we investigated the surface charge potential in more detail. As can be seen in the visualization in Figure 4c, variant 2 in particular contains a large positively charged surface region formed due to the introduction of the V99K/R mutation compared to the WT antibody. The positively charged surface patch is still present in the WT antibody but less pronounced, whereas the introduction of additional negative charges in variant 6 completely disrupts the patch (Figure 4c). This entails a drastic change in binding regime for variant 6 as additional repulsion makes additive electrostatic interactions unfavourable and, hence, no high molecular weight cluster formation and phase separation is observed.

To further characterize the phase behaviour of antibody–DNA condensates, and in particular map the chemical phase space in more detail, we applied PhaseScan^42^. This high-throughput droplet microfluidics-based approach allows for high resolution assessment of biomolecule phase diagrams, allowing us to further understand the nature of antibody–DNA interactions crucial to phase separation. We mapped out the phase space for the WT antibody and variant 2 over a broad range of conditions as shown in Figure 5a. Interestingly, the phase space is altered vastly between variant 2 and the WT antibody. Variant 2 phase separates less readily at lower DNA contents and more easily at higher DNA concentrations (Figure 5a, left panel). More specifically, a minimal DNA content of ∼ 4 µM is necessary to allow for phase separation of variant 2. Compared to that, the minimal DNA content necessary to allow for phase separation of the WT antibody is much lower and fell below the tested concentration range (Figure 5, right panel). The phase diagram at 0 µM DNA, however, confirms that the pure WT antibody does not undergo phase separation on its own. The difference in the necessary minimal DNA content for phase separation can be explained by the fact that variant 2 has a more pronounced positive charged patch. More specifically, this will allow the antibodies to interact more strongly with individual DNA strands and especially in large excess of antibody (*>* 10x) preferentiate formation of isolated DNA–antibody complexes. Hence, higher contents of DNA are required to introduce large scale bridging and ultimately phase separation. At higher DNA contents, such as 10 µM DNA, only ∼ 25 µM of variant 2 is needed to induce phase separation, whereas as much as ∼ 40 µM are necessary for the WT antibody. This is likely because the highly charged DNA can be stabilized more effectively by variant 2 as it exhibits an increased extent of the positive charged surface patch. Hence, in conditions where phase separation is observed, recruitment into condensates is expected to be more effective for variant 2 as compared to the WT antibody. Indeed, this correlates well with the above observation that variant 2 condensates immediately after formation are larger in size compared to the WT antibody shortly after formation. Hence, it becomes clear that the surface patch properties and changes therein, represented by the individual variants, govern and modulate the macroscopic phase separation observed.

**Figure 5:**
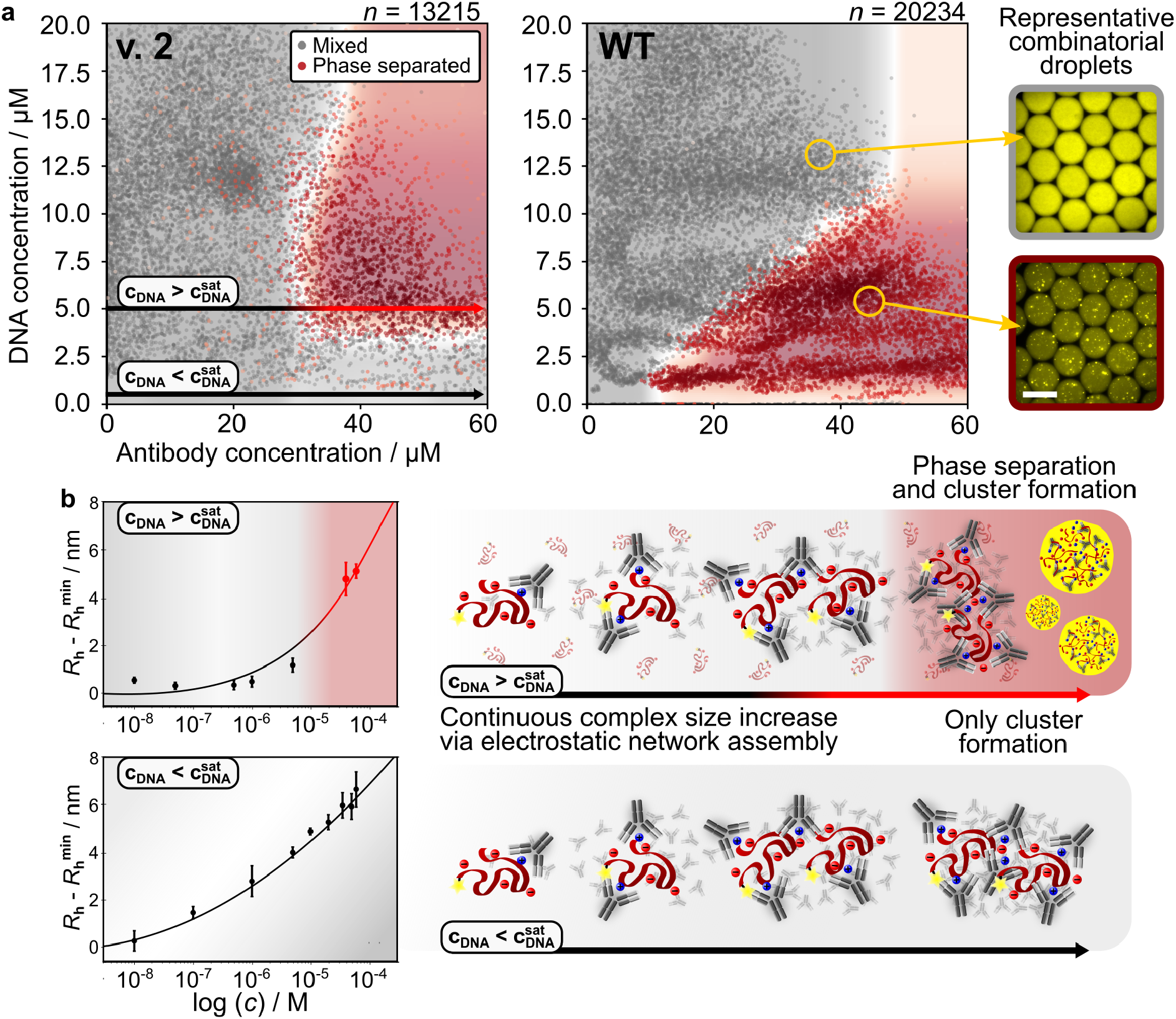
Mapping the chemical phase space and connecting it to high molecular weight cluster formation. **(a)** Phase diagrams of variant 2 and WT at room temperature as obtained by PhaseScan with grey points indicating a mixed phase and red points indicating phase separated conditions. At lower DNA contents, the WT antibody phase separates more readily whereas with increasing DNA contents variant 2 becomes more likely to undergo phase separation. Each point in the phase diagram represents a condition probed by a droplet generated with combinatorial microfluidics. (top) Droplets showing mixed phase and (bottom) phase separated conditions, scale bar represents 40 µm. **(b)** Binding curve titrations across horizontal cross sections (const. DNA, varying antibody concentration) of the phase diagram below (*c*_*DNA*_ = 0.1 µM) and above (*c*_*DNA*_ = 5 µM) the minimal DNA content necessary to undergo phase separation. Errors represent standard deviations from triplicate measurements. In both instances continuous increase of the complex size is observed. This indicates that network formation based on additive electrostatic interactions is an inherent property of the system, not just of phase separating conditions. Hence, the DNA saturation concentration functions as a critical concentration of available linkers between DNA–antibody clusters.

The detailed maps of the chemical phase space in combination with the microfluidics– based binding analysis further allows us to mechanistically rationalize the interplay between molecular binding events and macroscopic behaviour. More specifically, using MDS we studied the molecular binding events and cluster formation across horizontal cross sections (i.e., constant DNA, varying antibody concentration) of the phase diagram by tracing *R*^*DNA*^ (Figure 5a, b). To do so, we monitored DNA–antibody complex sizes with variant 2 antibody below (*c*_*DNA*_ = 0.1 µM, see Figure 5b, lower panel) and above (*c*_*DNA*_ = 5 µM, see Figure 5b, upper panel) the minimal DNA content necessary to undergo phase separation 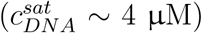 (Figure 5). At high DNA concentration conditions 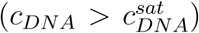, we observed a continuous complex size increase resulting from the formation of high molecular weight clusters with increasing antibody concentrations (Figure 5b). At high antibody concentrations (20, 40 and 60 µM) samples were then found to visibly become turbid and show phase separation. By separating the dense phase from the supernatant via centrifugation we were able to further characterise the size of DNA–antibody complexes in equilibrium with the dense phase. These complexes show further increase in their hydrodynamic radius along the initial trajectory even after phase separation was observed as is shown in Figure 5b, upper panel. Stunningly, at DNA concentrations where the system does not undergo phase separation (i.e.,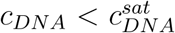) the DNA–antibody complexes also show a continuous increase in size and formation of high molecular weight clusters with increasing antibody concentrations (Figure 5b, lower panel). This suggests that electrostatic network assembly is an inherent property of the system, as opposed to an induced behaviour in phase separating conditions. Furthermore, it can be inferred that high antibody excesses are driving the formation of high molecular weight clusters, whereas the DNA content is crucial in inducing phase separation, as is schematically illustrated in Figure 5b.

## Discussion

In this work, we have presented a systematic study of the impact of surface patch property changes on antibody non-specificity using the HzATNP antibody library as a model system and DNA as a non-specificity ligand. In utilizing and establishing microfluidic diffusional sizing to probe non-specific interactions in the solution phase, we observed pronounced differences in the non-specific binding amongst different HzATNP antibody variants to DNA induced by only few mutations. Crucially, we have identified hydrogen bonding as the key determinant of non-specificity in the form of hydrogen bonding surface patches (i.e., clustering of residues able to engage in hydrogen bonding on the protein surface), which allow for concerted complexation to DNA strands. Disruption of this patch strongly decreased the non-specific binding affinities to DNA (Figure 6), hence, restored desirable behaviour. The WT antibody, for example, displays affinites of ∼ 1 µM, whereas the replacement of hydrogen bonds with charges or removal of hydrogen bonding capacity e.g. T68G decreases affinity by multiple orders of magnitude in variants 2, 4 and 6. Secondly, DNA induced liquid–liquid phase separation was discovered based on a change in condition to lowered ionic strengths (Figure 6). The decrease in Debye screening causes the electrostatic surface patch properties to become the dominating factor distinguishing between behaviours. More specifically, our experiments revealed continuous increase of DNA–antibody complex sizes with increasing antibody concentrations based on additive electrostatic interactions for a subset of the library. In tracing binding behaviours as well as the phase space for selected variants, we hypothesize that the extent of a positive charged patch on the protein surface can translate into a macroscopic behaviour. As such, variant 2 does phase separate, forming large condensates, while showing no binding saturation, whereas variant 6 does not phase separate and displays binding site saturation with increasing antibody concentration. Taken together, we observe changes in molecular binding events introduced by only individual mutations and connect them to the observed phase space to demonstrate the importance of formulation conditions to the balance between charged and hydrogen bonding surface patch properties and their impact on non-specificity binding and phase separation (Figure 6).

**Figure 6:**
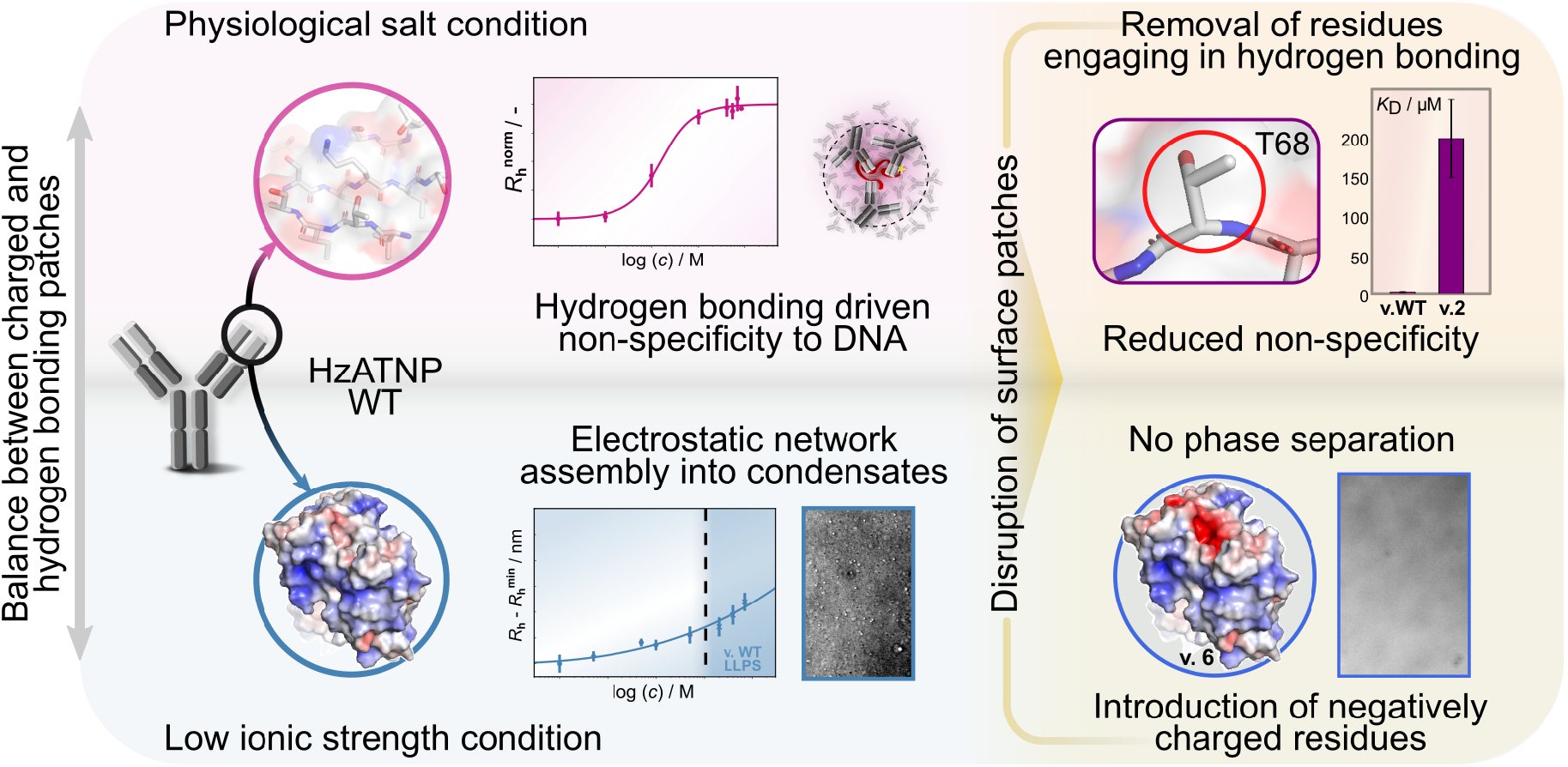
Balance between charged and hydrogen bonding patches governs non-specific binding and phase separation of antibodies with DNA. At physiological salt conditions, a patch of surface hydrogen bonding drives non-specific binding of the WT antibody to DNA. Disruption of the surface patch via mutation of critical residues drastically increases the dissociation constant and, hence, the non-specific binding behaviour. At lowered ionic strengths, the surface charge properties become dominant, revealing a connected positively charged region on the WT antibody surface. This leads to additive electrostatic interactions, which entail formation of high molecular weight clusters and liquid-liquid phase separation. Disruption of the positively charged region allows for disruption of the additive electrostatic interactions, which makes phase separation unfavourable. Taken together, this demonstrates an inherent balance of the impact of electrostatic and hydrogen bonding patches on phase separation and non-specific binding based on the formulation conditions.

Surface patches are gaining more and more attention in the field of protein developability as individual case studies identifying the molecular causes of non-specificity in detail commonly highlight these as key drivers. As such, patches of surface charge and hydrophobicity have already been shown to be able to vastly alter non-specific binding behaviour.^25,26,28,31,34^ The discovery of hydrogen bonding surface patches, as shown in this work, fits very well into this mould in representing another category of fundamental molecular forces common to protein binding interactions. Hence, hydrogen bonding patches are a crucial addition to identifying and modulating non-specificity by allowing for the explanation of various non-specific binding occurrences which are so far hard to resolve. Interestingly, patches of surface charge and hydrophobicity can usually be disrupted by introducing only individual mutations, a strategy that also has been found successful here for the case of the HzATNP library. Furthermore, the ability of such hydrogen bonding patches to drive non-specific binding to DNA, could be crucially important, as DNA non-specific binding has been shown to correlate well with non-specific binding propensity *in vivo*^27^. As such hydrogen bonding patches could be a key contributor to problematic pharmacokinetics and *in vivo* behaviour, which further highlights the importance of considerations of the surface patch properties to studying non-specificity and developability overall.

Antibody phase separation has been described previously for single component systems, where strong surface charge asymmetries generate regions with complementary charges on an individual antibody.^47–49^ In these systems, additive electrostatic interactions typically drive formation of antibody networks and eventually nucleate condensates or form gels. Interestingly, the system studied here exhibits similar features in that additive electrostatic interactions are present. The complementary charges, however, are split onto two components in antibody and DNA. Notably, it is well established that nucleic acids such as DNA/RNA allow for complex coacervate and condensate formation, particularly with peptides and proteins^50,51^. To the best of our knowledge, this represents the first description of DNA-induced phase separation of antibodies. This potentially represents a key finding to the tendency of larger, folded proteins to undergo phase separation and considerations for future *in vivo* activity and formulation processes of therapeutic proteins. This reveals that based on the formulation conditions there is a inherent balance between molecular properties that can vastly change behaviours. Lastly, the binding analysis presented in this work, provides a valuable example that the study of the molecular binding processes can mechanistically predict macroscopic behaviours such as phase separation. Therefore, we suggest that the interplay of specific binding and interaction network formation may be a crucial parameter for understanding the molecular basis of heterotypic phase separation.

Non-specificity has long been an elusive topic to grasp, with the responsible molecular features being largely unclear. Investigations have featured sequence based net properties^52–54^ or specific abundances of individual amino acids^55–59^. These approaches, however, often fail to impact non-specificity directly and selection guidelines are largely limited to excluding candidates with properties to the extremata of common distributions. In showing the vast impact of only small variations on the surface patch properties such as the hydrogen bonding propensity, our study gives clear indication that these are the key molecular features altering protein colloidal properties overall and with that non-specificity. Crucially, only individual mutations within a 150kDa antibody protein are sufficient to alter non-specific binding affinities drastically by orders of magnitude or entirely reverse a macroscopic behaviour in phase separation of antibody–DNA samples. Overall, this indicates the potential of investigating surface patch properties to optimise future development pipelines. More specifically, this could eventually allow for a potential reversal of the development process from searching for causative properties after identification of problematic behaviour to identifying property profiles and anticipating undesired issues.

## Materials & methods

### HzATNP antibody variants

Design, characterisation and expression of the HzATNP antibodies have been performed as described in detail previously^30,37^. After expression, all antibody samples were stored in aliquots at roughly 5-10 mg/mL in either 20mM HEPES buffer pH = 7.4, 150mM NaCl, with 0.01 % (*v/v*) Tween (HS buffer) or 2mM HEPES buffer pH = 7.4, 15mM NaCl, with 0.01 % (*v/v*) Tween (LS buffer) at −80°C. Protein stocks were thawed on ice and shock frozen using liquid nitrogen, up to a maximum of three freeze-thaw cycles.

### DNA and RNA oligos

DNA oligos were purchased from Merck (Darmstadt, Germany; HPLC purified and dry). Strands included, a 5’-modification with Cy3. The 100 mer, 50 mer and 20 mer sequences used were CTCACCCACAACCACAAACAATTTAAATAATATTAAATAATA-TTAATATATTATCGATTAAATAATAATTAATTAATATTGGTTGGATGGTAGATGGTGA, CTCACCCACAACCACAAACAATTTAAATAATATTAAATAATATTAATATA and CT-CACCCACAACCACAAACA, respectively. DNA oligos were then dissolved in double deionized (dd) water or 2 mM HEPES buffer pH = 7.4, 15mM NaCl, with 0.01 % (*v/v*) Tween 20. PolyA RNA lyophilized was purchased from Merck (Darmstadt, Germany). PolyT RNA 20 mer, including a 5’ modification with Cy5, was purchased from Biomers (Ulm, Germany)

### Fabrication of microfluidic diffusional sizing devices

Fabrication of microfluidic diffusional sizing (MDS) devices was performed according to previous descriptions^39^. First, devices were designed using AutoCAD software (Autodesk, San Rafael, CA, USA), followed by printing of the designs onto acetate masks (Micro Lithography Services, Essex, UK). These were used to generate SU-8 3025 molds to result in approx. 25 µm device heights utilising photolithographic methods. To do so, silicon wavers were spin coated with SU-8 photo resist at 500 rpm for 10 seconds followed by 3000 rpm for 45 seconds and subsequently baked for 15 minutes at 95°C. Acetate masks were then placed on the coated waver and exposed to UV light at 365 nm for 40 seconds, followed by heating at 95°C for 5 minutes. The SU-8 mold was then developed by washing with PGMEA for 10 min under vigorous shaking and rinsing with IPA. Device heights were then determined using a profilometer (Dektak, Bruker, USA). The master was then placed in a Petri-dish which was filled with PDMS (2 component system, Momentive, Techsil, Bidford-on-Avon, UK) supplemented with carbon black nanoparticles. The black PDMS was subsequently degassed in a desiccator until complete removal of bubbles (approx. 2h) and afterwards baked for 1h at 65°C. After Baking the PDMS mold was removed from the silicon waver, cut into individual chips as well as punched to yield inlet and outlets, all under constant removal of dust particles. The processed PDMS devices were then sonicated for 15 min in IPA and the glass/quartz slides in EtOH. After sonication both PDMS devices and cover slips were dried under air flow and subsequently surface radicalised in an oxygen plasma oven (Femtro from Electronic Diener, Ebhausen, Germany) at a plasma power of 40% for 30 seconds. Cover slips and black PDMS devices are then bonded together by simple contacting both treated surfaces followed by baking for 10 min at 65°C.

### Microfluidic diffusional sizing of DNA and DNA–HzATNP antibody complexes

A schematic illustration of the chip design is shown in Fig. 1. Operation of the MDS devices was performed according to previous descriptions.^39,40^ In brief, microfluidic devices were filled with experiment buffer, in this case 20 mM HEPES buffer pH = 7.4, 150 mM NaCl, with 0.01 % (*v/v*) Tween or 2 mM HEPES buffer pH = 7.4, 15 mM NaCl, with 0.01 % (*v/v*) Tween. Negative pressure was applied to the outlet via a glass syringe (Hamilton, Bonaduz, Switzerland) connected to a syringe pump (neMESYS, Cetoni GmbH, Korbussen, Germany), which allowed for control of flow rates and sample loading through reservoirs connected to the respective inlets. The sample inlet reservoir was provided with the sample conditions specified for each data set and the co-flow buffer inlet reservoir was provided with the matching buffer i.e. either 20 mM HEPES (pH 7.4), 150 mM NaCl, 0.01% Tween or 2 mM HEPES (pH 7.4), 15 mM NaCl, 0.01% Tween. Samples were kept and incubated in specified temperature conditions thorughout the experiment. Images were recorded through a custom-built inverted epifluorescence microscope which was equipped with a CMOS camera (Prime 95B, Photometrics, Tucson, AZ, USA), brightfield LED light sources (Thorlabs, Newton, NJ, USA) and a Cy3-laserline filter cube (Laser2000, Huntingdon, UK). Images were taken at flow rates of 60 *µ*L/h and 100 *µ*L/h, with three individual repeats each. Extraction of the diffusion profiles was performed via analysis of the fluorescence images with a custom written analysis software.^53^ After popagation of the intitial profiles with analytical model numerical model simulations are fitted, which are solutions to the diffusion–advection equations for mass transport under flow^40^.

### Diffusional sizing of HzATNP antibodies via intrinsic fluorescence

Intrinsic fluorescence was used to detect HzATNP antibody diffusion profiles and was only applied in high excesses of antibody, where the intrinsic fluorescence signal of DNA could be neglected. For UV measurements, quartz instead of glass cover slips were used to mitigate the absorption of glass itself. Here, a custom-built inverted microscope which was equipped with a 280nm LED light source (M280L3, Thorlabs, Newton, NJ, USA), a charge-coupled-device camera (Prime 95B, Photometrics, Tucson, AZ, USA) and filter set Semrock TRP-A-000 (Laser2000, Huntingdon, UK) was utilized as described previously.^53,60^

### DNA–HzATNP antibody library phase separation

Phase separation was exclusively induced in vitro and specific conditions are reported for each specific dataset. Generally, phase separation was induced by gently mixing the individual components in 500µL centrifuge tubes. Buffer (2 mM HEPES buffer pH = 7.4, 15 mM NaCl, with 0.01 % (*v/v*) Tween) was added first, followed by DNA and then antibody. Usually 10 *u*L of 40 µM antibody and 5 µM DNA 50 mer were prepared. Images were obtained by application of 1-2 µL of sample onto a cover slip, followed by immediate imaging in sealed chambers. For phase separation experiments of the HzATNP library with PolyA, PolyA was spiked with 5 % (*w/v*) of PolyT Cy5 labelled, to give a stock solution of 15 mg/mL in 50 mM HEPES buffer pH = 7.2, which was diluted in the sample to yield buffer conditions of 2 mM HEPES (pH 7.4), 15 mM NaCl, 0.01% (*v/v*) Tween. Specific conditions for the usage of 20 mer, 100 mer and PolyA are described in detail where shown. Fluorescence was observed using an inverted microscope (OpenFrame, CairnResearch, Faversham, UK) equipped with a 20x air objective (Nikon, Surbitton, UK) and appropriate filters (Laser2000, Huntingdon, UK). All images were collected with a high sensitivity camera (Prime BSI Express sCMOS, Photometrics, Tucson, AZ, USA).

### Dissolution experiments

For the dissolution experiments DNA–HzATNP antibody phase separating samples at 50 µM variant 4 and 6.25µM DNA 100 mer were prepared as discussed previously. Then 20 *v/v* % of 1M NaCl or 50 % (*w/v*) 1,6-hexanediol were added to give sample conditions of 40 µM variant 4 and 5 µM DNA 50 mer plus 212 mM sodium chloride or 10 % (*w/v*) 1,6-hexanediol, respectively. Imaging was performed as reported above.

0 *µ*M antibody, obtained by addition of 20 % (*v/v*) of either 1M NaCl leading to 212 *µ*M NaCl or of 50 % (*w/v*) 1,6 hexanediol leading to 10 % (*w/v*) 1,6 hexanediol.

### Mapping chemical phase space using PhaseScan

PhaseScan was operated according to previously described procedures^42^. Briefly, droplets were generated on the microfluidic device using automated syringe pumps. This allows for combination of the aqueous droplet components prior to the droplet-generating junction. Through variation of the flow rates, the combination of droplet composition can be varied. More specifically, the three input streams were: buffer (2mM HEPES buffer, pH = 7.4, 15mM NaCl), 16 µM Cy3 labelled DNA 50 mer in buffer supplemented with 4 µM Atto 488 dye, and 75 µM HzATNP antibody of specified variant in buffer supplemented with 4 µM Alexa 647 dye. For the generation of microdroplets, FC-40 oil (containing 1% (*w/v*) fluorosurfactant, RAN biotechnologies) was flown into the chip at 150 µL/h of constant flow rate. Droplets were then imaged subsequently with an inverted microscope (OpenFrame, Cairn Research, Faversham, UK) using a 20x objective and collected through a camera (Kinetix, Photometrics, Tucson, AZ, USA). Appropriate filters were applied to allow for fluorescence detection. Based on the fluorescence intensity in each channel, every droplet could be mapped to a unique point in phase space and assessed for phase separation.

### Homology model of HzATNP variants

The homology model of the WT antibody was obtained as previously described using Modeller (version 9.12).^30^ Structural models of all other HzATNP variants were subsequently created through mutagenesis of diverging residues from the WT antibody via PyMOL2. Vacuum electrostatic surface potentials were generated using the ABPS Electrostatics PyMOL2 plug-in on the existing models.

## Author Contributions

H.A., T.W.H., G.K. and T.P.J.K. designed and conceptualized the study. H.A., T.J.W. and T.S. performed experiments. N.L.Z., G.I., T.W.H., G.K., T.S., and T.P.J.K. provided materials and methods. H.A., M.M.S, G.K. and T.P.J.K. analysed the data. H.A. and G.K. wrote the original draft of the paper. All authors discussed the results and commented on the manuscript.

## Acknowledgements

The research leading to these results has received funding from Global Research Technologies, Novo Nordisk A/S (H.A., T.H.W., T.P.J.K.), the European Research Council under the European Union’s Horizon 2020 Framework Programme through the Marie Sklodowska-Curie grant MicroSPARK (agreement n 841466; G.K.), the Herchel Smith Funds (G.K.), and the Wolfson College Junior Research Fellowship (G.K.). TJW thanks the Harding Distinguished Postgraduate Scholar Programme. We thank Dr. Thomas Egebjerg and Dr. Jais Rose Bjelke from Global Research Technologies, Novo Nordisk A/S for their help with generation of the antibody library. We thank Dr. Thomas Egebjerg and Dr. Jais Rose Bjelke from Novo Nordisk A/S for their help with generation of the antibody model system.

## Data availability

The raw data and analysis code underlying this study will be made available upon reasonable request.

