## Supplementary Information for "Surface interaction patches link non-specific binding and phase separation of antibodies"

#### Affinity and stoichiometry analysis

To asses the affinity of the DNA–HzATNP antibody binding complexes, we first consider the reaction between DNA binding site $DNA_b$ and antibody binding site $mAb_b$ forming binding site complex C:

$$DNA_{\mathrm{b}} + mAb_{\mathrm{b}} \rightleftharpoons C$$

The association equilibrium is then given by

$$K_{\mathrm{a}} = K_{\mathrm{d}}^{-1} = \frac{[C]}{[DNA_{\mathrm{b}}][mAb_{\mathrm{b}}]} \tag{1}$$

where $K_{\mathrm{d}}^{-1}$ is the dissociation constant. The following material balances hold with respect to this binding equilibrium:

$$[DNA_{\mathrm{b}}]_0 = [DNA_{\mathrm{b}}] + [C] \tag{2}$$

$$[mAb_{\mathrm{b}}]_0 = [mAb_{\mathrm{b}}] + [C] \tag{3}$$

After insertion of the material balances in the binding equilibrium one can solve for [C]. Assuming there is no cooperativity in complex binding the total number of binding sites can be replaced with the number of individual binding sites to correct for the stoichiometry of the binding reaction:

$$[DNA_{\mathrm{b}}] = n_{DNA} * [DNA]$$

$$[mAb_{\mathrm{b}}] = n_{mAb} * [mAb]$$

Picking the only physically possible solution for C then yields:

$$[C] = \frac{n_{DNA} * [DNA]_0 + n_{mA} * [mAb]_0 + K_{\mathrm{d}}}{2} - \sqrt{\left(\frac{n_{DNA} * [DNA]_0 + n_{mAb} * mAb]_0 + K_d}{2}\right)^2 - n_{DNA} * n_{mAb} * [DNA]_0 * [mAb]_0}$$

Pertaining knowledge of the complex stoichiometry the concentrations can then be related to the observed hydrodynamic radius using the following assumptive relation to allow for determination of the affinity:

$$R_{\mathrm{h}}^{Obs} = R_{\mathrm{h}}^{0} + \Delta R_{\mathrm{h}}^{Max} \frac{[C]}{[DNA_{\mathrm{b}}]} \tag{4}$$

Here $R_{\mathrm{h}}^{0}$ is the lower plateau value and $\Delta R_{\mathrm{h}}^{Max} = R_{\mathrm{h}}^{Max} - R_{\mathrm{h}}^{0}$ the difference between both plateau values. The binding complex stoichiometry was determined via performing binding curve analysis of DNA–WT antibody complexes at 50, 1000 and 2000 nM DNA concentration see Figure. This is necessary to allow for constraining of the affinity/stoichiometry plane. This yielded $n_{DNA} \approx 1$ and $n_{mAb} \approx 2.7$, which was further utilized also for DNA binding complexes with all other variants. This is also consistent with the hydrodynamic radius of 6.5 nm largely similar for all variants and conditions showing binding site saturation, which according to common prediction tools corresponds to a molecular weight of approx. 500 kDa assuming a globular complex. Based on the stoichiometric analysis the antibodies would account for 405 kDa (150 kDa x 2.7) whereas a theoretical folded weight of 70 kDa can be inferred from the pure DNA oligo (3.7 nm), giving a total of 475 kDa. Similarly for a complex of 10 nm in hydrodynamic radius, as found for variants 1, 2 and 4 and lowered ionic strengths and 1 $\mu$M DNA and 6.7 $\mu$M antibody, a total weight of $\sim$ 1700 kDa can be inferred. This, when assuming recruitment of additional DNA and antibody into the complex with the same stoichiometry as found previously, would entail $\sim$ 10 antibodies and $\sim$ 3 DNA strands to be bound in one complex.

### Supplementary figures

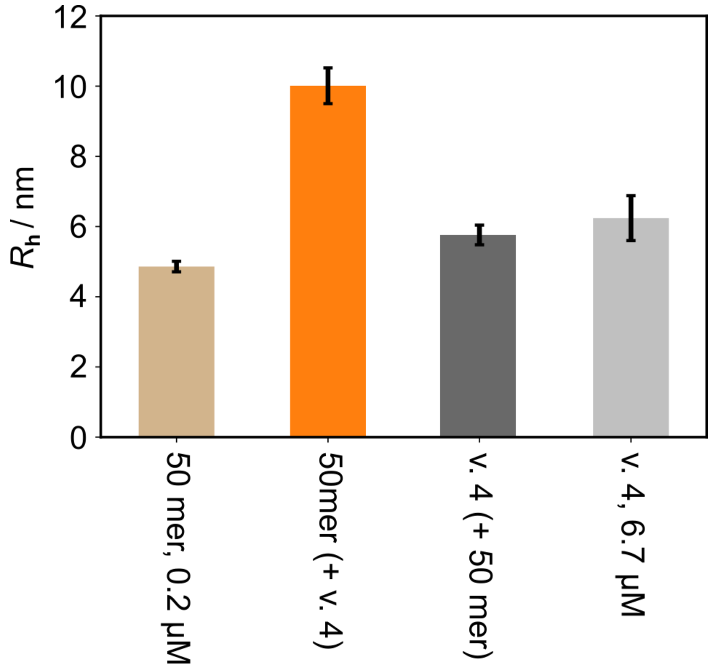


**Supplementary Figure S1:** Mircofluidic diffusional sizing of DNA 50 mer (left bar), DNA 50 mer–HzATNP variant 4 antibody (middle two bars) and HzATNP variant 4 antibody (right bar) under lowered ionic strength conditions (2 mM HEPES buffer pH = 7.4, 15 mM NaCl). DNA signals are recorded using fluorescence microscopic of the Cy3 labelled DNA and antibody signal is recorded using intrinsic fluorescence of the protein assuming significantly higher signal from the antibody given a high excess. The individual components in DNA and antibody are not causing the high molecular weight cluster formation as the clusters are only formed in the presence of both. Given the high excess of antibody both antibody in the mixture and alone appear to have the same size as the majority of the signal obtained from the mixture is from free antibody.

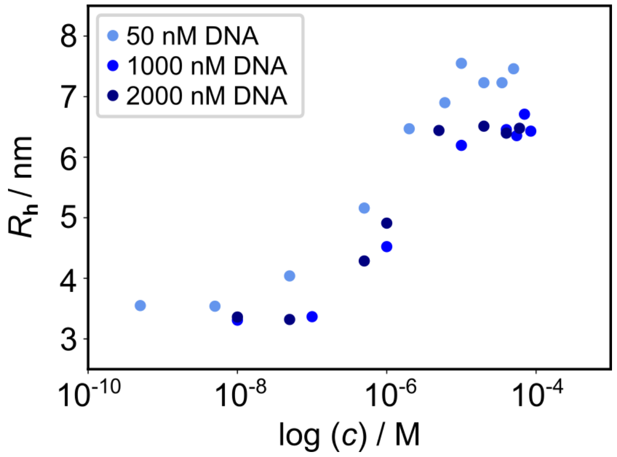


**Supplementary Figure S2:** Binding curves of WT antibody at multiple DNA concentrations (50, 1000 or 2000 nM DNA). This is used to perform stoichiometric analysis by constraining Eq. 4 not only with respect to $K_D$ but also $n_{DNA}$ to determine the number of antibody binding sites on an individual DNA strand.

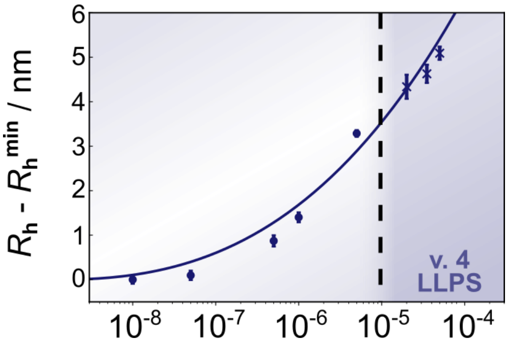


**Supplementary Figure S3:** Binding curve titration of variant 4 antibody against DNA under lowered ionic strength conditions (20 mM HEPES buffer, pH = 7.4; 150 mM NaCl, 4 $°C$). Complex sizes display continuous increase with increasing antibody concentration.

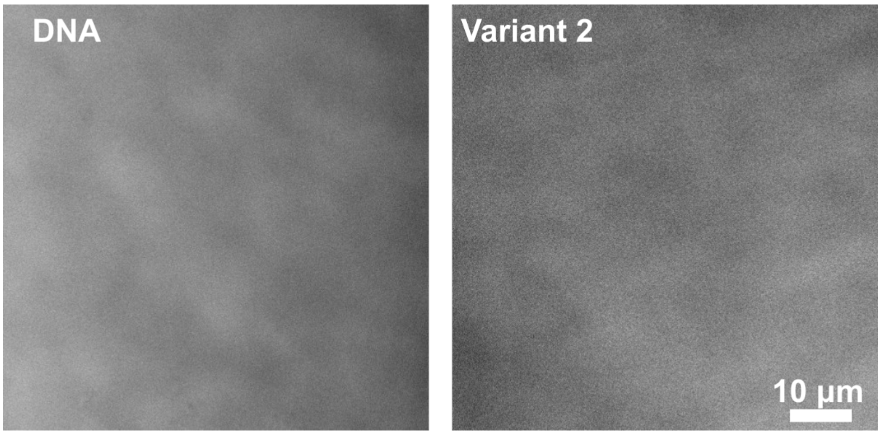


**Supplementary Figure S4:** DNA 50 mer (left, 50 $\mu$M) and variant 2 antibody (right, 50 $\mu$M) do not undergo phase separation individually. All images are brightfield images.

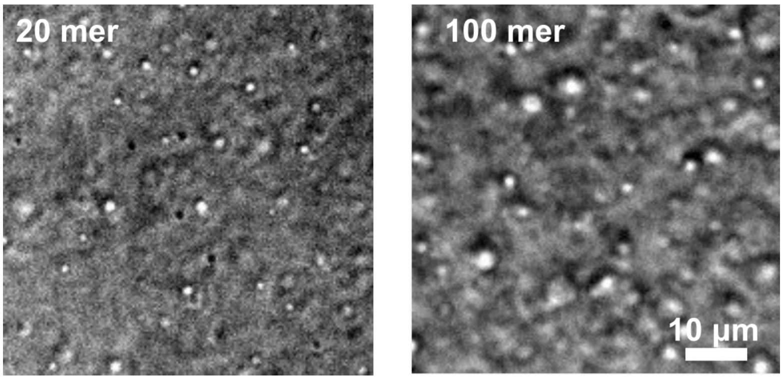


**Supplementary Figure S5:** 20 mer (left) and 100 mer (right) DNA strands also undergo LLPS with HzATNP variant 4 antibody. Condensates were formed at 5 $\mu$M DNA 20 mer and 60 $\mu$M antibody and 2 $\mu$M DNA 100 mer and 20 $\mu$M antibody, respectively. All images are brightfield images.

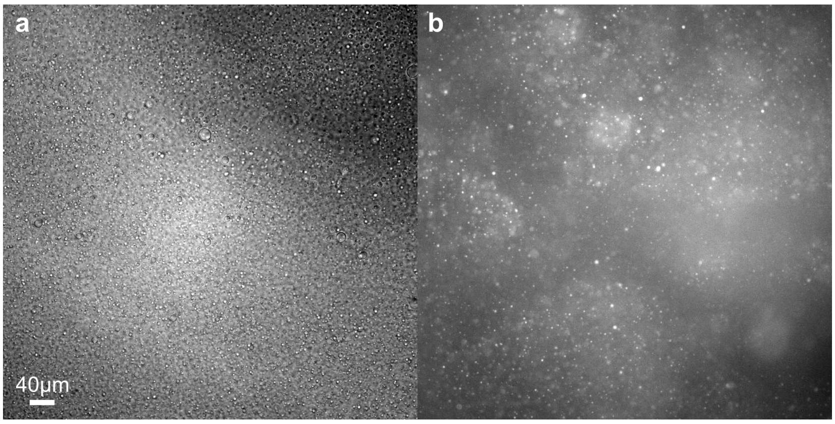


**Supplementary Figure S6:** Brightfield **(a)** and A647 fluorescence **(b)** imaging of PolyA–HzATNP v. 4 condensates at 6 $\mu$M antibody and 270 ng/$\mu$L of PolyA RNA.

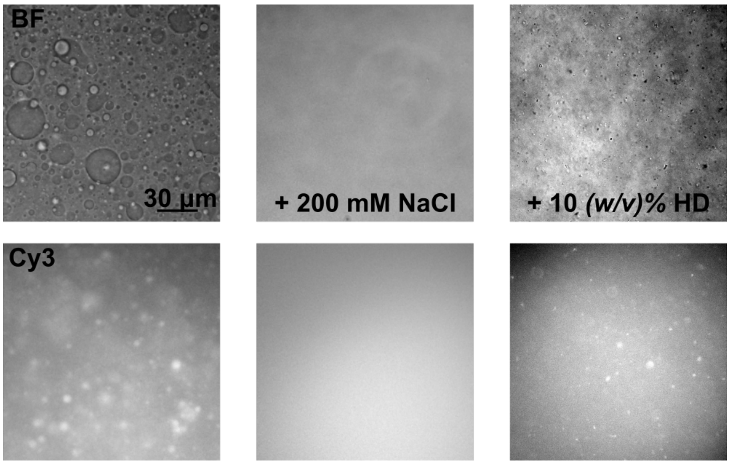


**Supplementary Figure S7:** Condensates formed at 6.25 $\mu$M DNA 50 mer and 50 $\mu$M HzATNP v. 4 (left) can be dissolved by introduction of high salt content (middle). Upon addition of a hydrophobicity disruptor in 1,6-hexandiol condensates are also dissolved but formation of aggregates is observed. Final conditions are 5 $\mu$M DNA and 40 $\mu$M antibody, obtained by addition of 20 % (*v/v*) of either 1M NaCl leading to 212 $\mu$M NaCl or of 50 % (*w/v*) 1,6-hexanediol leading to 10 % (*w/v*) 1,6-hexanediol.
